# The Evolution and Epidemiology of H3N2 Canine Influenza Virus After 20 Years in Dogs

**DOI:** 10.1101/2024.07.19.604289

**Authors:** Brian R. Wasik, Lambodhar Damodaran, Maria A. Maltepes, Ian E.H. Voorhees, Christian M. Leutenegger, Sandra Newbury, Louise H. Moncla, Benjamin D. Dalziel, Laura B. Goodman, Colin R. Parrish

## Abstract

The H3N2 canine influenza virus (CIV) emerged from an avian reservoir in Asia around 2004. As the virus has now been circulating entirely among dogs for 20 years, we here update our understanding of the evolution of virus in its new host. As a host-switched virus, H3N2 CIV will also reveal any host-adaptive changes arising during thousands of infections within its new host, and our analysis showed that the virus has evolved at a constant rate. CIV was first introduced into North America in 2015 from Korea, and we specifically examined the epidemiology of the virus among dogs in North America since then, including local outbreaks, regional die-outs, and repeated reintroduction from Asia. The H3N2 CIV now appears endemic only in China after dying out in South Korea around 2017. Virus lineages circulating in China appear to have seeded the most recent US outbreaks – with 2 or 3 introductions into North America during the past 3 years. Combining clinical reports, diagnostic testing data, and analysis of viral genomes we show that the virus spreads rapidly among dogs in kennels and shelters in different regions – likely dying out locally after all those animals become infected and immune. The overall epidemic therefore requires longer-distance dispersal of virus to initiate outbreaks in new locations. Patterns of spread in the USA may select viruses most adapted to those dense populations, which may lack the properties required for efficient long-distance transfers to other dog populations that would keep the virus in prolonged circulation.

**IMPORTANCE:** Viruses occasionally jump into new hosts to cause epidemics and may spread widely due to movement of humans or animals, or their viruses, with profound consequences for global health. The emergence and epidemiology of new epidemic viruses in companion animals provides a model for understanding disease dynamics and evolution. The H3N2 canine influenza virus arose from an avian virus, and infected dogs provide many opportunities for human exposure. H3N2 CIV transmission is dominated by fast-moving outbreaks within dense populations in animal shelters or kennels, while sustaining the epidemic likely requires movement of virus to more distant dog populations. Viral spread within North Americahas only been sustained for a few years at a time after which the virus dies out. The epidemiological and evolutionary dynamics of this virus in this structured host population shows how an acute respiratory pathogen can emerge and spread in a new host and population.

## Introduction

Influenza A viruses (IAV) are extremely successful viral pathogens of vertebrates that are maintained within many natural bird reservoir populations that cause multiple spillovers and outbreaks in mammalian hosts [1]. IAV is an enveloped virus of the family *Orthyomyxoviridae*, with a negative-sense genome arranged into eight units: PB2, PB1, PA, HA, NP, NA, M, and NS [2]. Some IAVs have overcome a variety of different host barriers to emerge as epidemic or pandemic pathogens in humans and domesticated or wild animals, including swine, horses, dogs, seals, cats, and mink [1,3–5]. Spillovers of avian-origin IAV also frequently result in acute disease in other mammals or domestic poultry with little or no onward transmission [6]. Within the human population, the IAV subtype H1N1 emerged around 1918, and that later underwent reassortment with additional avian strains to create the H2N2 and H3N2 subtypes [2]. The emergence of IAVs to cause epidemics in new animal hosts provides insights into the processes and principles of viral host-switching, allowing us to better understand potential human pandemic viral emergence.

Carnivore mammals (members of the Order Carnivora), including dogs, have long been identified as being susceptible to infection by influenza A viruses [3,4]. This has included the observation of canine infection with seasonal human influenza strains [7–9], in addition to the successful experimental infection of dogs with a human H3N2 subtype [10]. The first sustained outbreak of an emergent canine influenza virus (CIV) occurred around 1999 after the transfer of an equine influenza subtype H3N8 to dogs. That outbreak in the United States of America (US) was only recognized in 2004 [11], spreading through much of the US and persisting until 2016 [12]. The epidemiology of H3N8 CIV was largely driven by transmission within and among shelters and kennels with high population turnover [13], such that viral lineages were strongly geographically clustered with outbreaks in major metropolitan areas, where the virus mostly died out in those areas in only a few months [14]. The story of H3N8 CIV suggested that dog population structure (at least in the US) was not ideal for influenza to develop into a sustained epidemic or endemic infection [12].

A lineage of avian-derived H3N2 emerged into dogs in eastern Asia (mainland China or Korean peninsula) around 2004, resulting in sustained circulation that persists to present day [15–17]. Viral sequence analysis revealed that the population quickly became separated into several geographic subclades in Asia, showing that the virus had been circulating separately in a number of different geographic regions [16]. In early 2015 the H3N2 CIV was first identified in the North American continent as the cause of an outbreak in the US around Chicago, Illinois [18]. That virus was initially introduced from Korea, and additional outbreaks occurred in many parts of the US (including around Chicago and in Georgia, Alabama, and North Carolina) but were largely controlled during early 2017 [19]. Subsequent H3N2 CIV outbreaks occurring later in 2017 among dogs in the US were identified with a temporal gap from earlier cases and impacting independent locations, including among states in the Midwest (Minnesota, Ohio, Indiana, Kentucky) and Southeast (Florida, Georgia, North Carolina, South Carolina) [19,20]. In 2018 further unique outbreaks of H3N2 CIV were identified in Ontario (Canada), without clear sourcing from the neighboring US [20,21]. Based on the geographic and temporal spread of these latter cases, as well as on the analysis of viral sequences, it was apparent that multiple international introductions of the H3N2 virus were involved in sparking the North American outbreaks that were seen between mid-2017 and 2018.

Viral emergence events in new hosts start with single infections resulting from spillovers, but those rarely go on to cause outbreaks or epidemics [22]. The emergence of viruses in epidemic forms is generally thought to be associated with the acquisition of host-adaptive mutations that allow better replication of the virus in the cells and tissues of the new host animal, as well as causing increased transmission [23]. These host-adaptive mutations are expected arise quickly after transfer to the new host – either during the first rounds of replication after spillover, or during the first animal-to-animal transfers [24–26]. While early selection of such mutations can be obvious shortly after the bottleneck that occurs upon transfer to the new host, later virus populations will diverge by sequence within the new host; thus, without strong molecular data sets, key adaptive changes are harder to define in the background of mutations arising from genetic drift and/or other selection pressures, including the increasing levels of host immunity [27–29].

Other factors associated with emergence of respiratory viruses include their epidemiology and host ecology. These include the virological influences of incubation times and shedding patterns, as well as host population structures – such as heterogeneity in the host density and distribution, as well as movement or other connections between separated populations [30–32]. The human population is dense, well-connected, and globally mobile, resulting in few ecological barriers to global transmission of endemic and newly emerged respiratory pathogens, as shown with seasonal influenza dynamics and during the rapid global spread of the H1N1 pandemic in 2009, or SARS CoV-2 in 2020 [33–36]. Other animals live in varying size groups, including smaller and isolated populations in the wild where pathogen transmission is self-limiting, or in large flocks or herds that allow efficient transfer at least within that population, while some animals have long range movements including globe-spanning migrations [1,37,38]. Domestication and farming of animals can result in animals gathered into novel structures: large well-connected populations on farms, dense animal markets, and other forms of human-directed movements that allow new modes of spread and maintenance of pathogens compared to those seen in the wild populations of the same hosts [39,40].

Companion animals such as dogs and cats have structured populations with additional features compared to those of domestic livestock. Those populations include many households with one or a few animals, as well as large and well-connected populations within kennels or animal shelters. Additional populations in some regions of the world include street animals that likely have variable levels of density and connectivity. In Africa, these complex population structures of dogs and anthropogenic effects of nearby human populations impact Rabies virus (RABV) dispersal, transmission, and evolutionary dynamics [41,42]. In the United States, dogs and cats live in close proximity to humans, so infected dogs would results in high levels of exposure to humans of all ages and health status [43].

Here we continue to examine the emergence, evolution, and epidemiology of H3N2 canine influenza A virus that has been circulating in its new host for 20 years, and provide a detailed understanding of the processes that allowed its emergence and sustained transmission. While the focus is on the epidemiology and evolution of the virus over the most recent 5 years in the USA, those are compared to those seen in during the previous 15 years. Recent cases in North America appear to have originated from Asia, with introductions occurring between 2017 and 2019, and again in 2021 and 2022, resulting in the current observed lineages.

## Results

### The global H3N2 CIV population initiated around 2004, has transferred repeatedly around Asia and North America, and formed multiple clades

We assembled a complete phylogenetic analysis of full H3N2 CIV genomes since 2006 (**Fig. 1**). Our data set combines obtained sequences for this study with others in public repositories to generate a large-scale H3N2 genome data (n = 297, **Table S1**) that revealed the global virus evolution in dogs. Initial phylogenetic analysis of each individual gene segment gave concordant results (**Fig. S1**), indicating that lineages did not result from extensive reassortment of the viral gene segments or exchange with other viral strains – which was further confirmed by reassortment analysis (**Fig. S2**). Full-genome alignments and phylogenetic analysis of H3N2 CIV reveals that virus population lineages can be classified into a series of distinct clades, which allow us to track circulation, population variants, and interconnectedness during different stages of the overall epidemic (**Fig. 1**).

**Figure 1.**
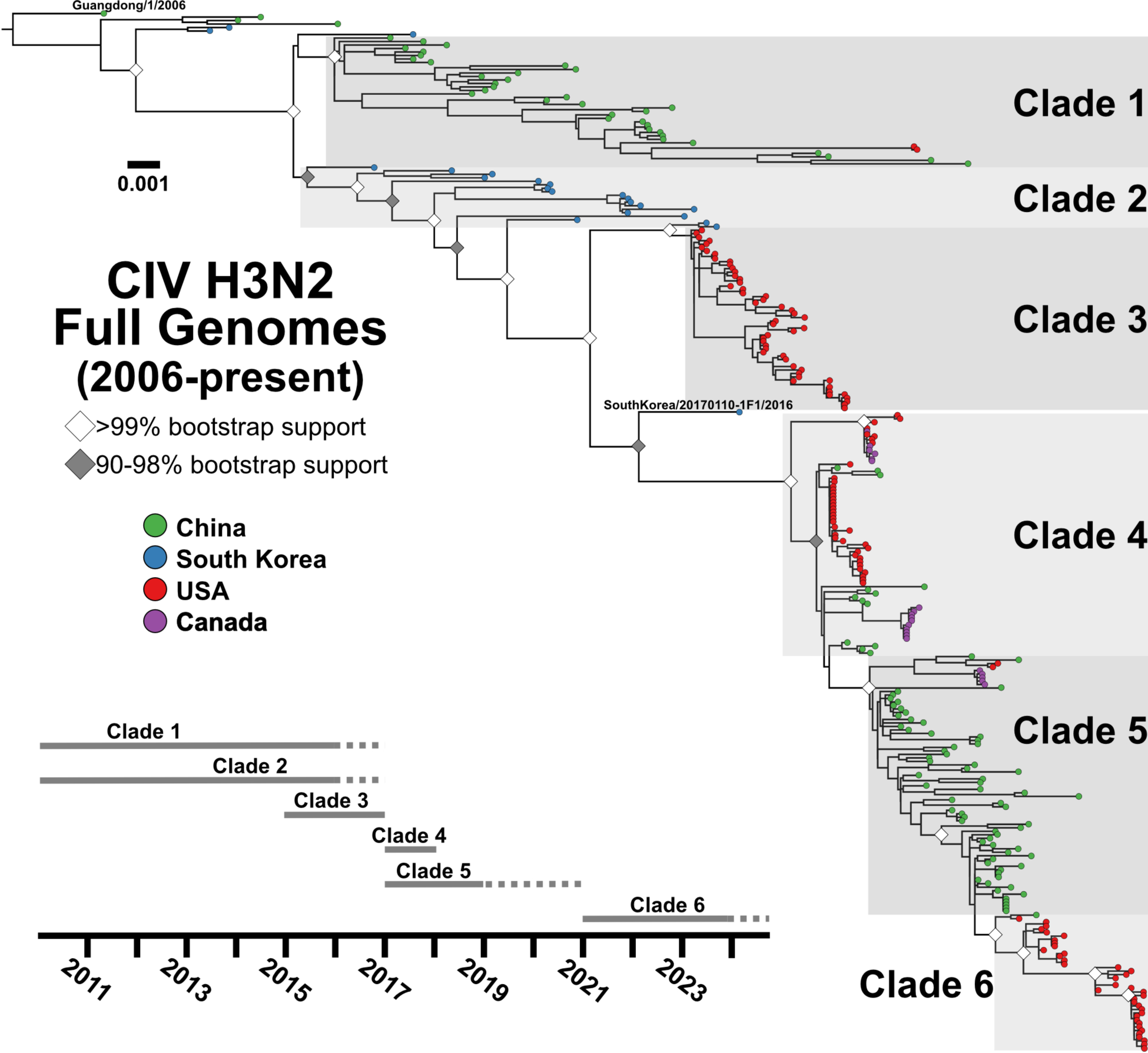
Overall global clade structure of H3N2 CIV genetic diversity. ML tree of the full genome data set (n=297) revealing distinct early lineages and corresponding clades (shaded and labeled). Tips denote geographic sampling source (green = China, blue = Korea, red = US, purple = Canada). The phylogeny is rooted on the sequence Guangdong/1/2006. Scale denotes nucleotide divergence. White diamonds at nodes represent bootstrap support >99%, grey is support from 90-98%. A general timeline of observed sampling of each clade is presented. The final Korean isolate linking Clades 2 and 4, SouthKorea/20170110-1F1/2016, is noted.

Emergent H3N2 CIV prior to 2010 reveal extensive transmission of viruses within Asia, primarily within and between China and South Korea [15,17], along with a reported limited outbreak in Thailand [44], until more geographically distinct lineages were established in China (Clade 1) and South Korea (Clade 2). Surveys in South Korea reported fewer cases of CIV between 2016 and 2018, with the Clade 2 virus lineage likely dying out in South Korea around that time [45,46]. Clade 1 viruses were infrequently reported in China after 2015, while CIV surveillance samples from dogs collected from 2017-2019 generated viral sequences that appeared to more resemble a Clade 2 derived virus that had first been circulating in South Korea. As such, a new clade (Clade 5) became established in China across several provinces, likely displacing the earlier viral lineages (Clade 1) [47–49].

Clades and lineages of North America were not derived until after 2015 and show connections to several clades originating in Asia in that time (**Fig. 1**). The first recorded introduction of H3N2 CIV into North America in early 2015 caused a series of outbreaks in the vicinity of Chicago, Illinois, and in several southeastern US states, causing a scattered set of outbreaks among dogs through early 2017 [19]. Phylogenetic analysis of the viral sequences showed that the virus in US dogs was derived from a Clade 2 virus circulating in South Korea, and that US lineage is herein now referred to as Clade 3 (**Fig. 1**). Our previous study also noted that later in 2017, US outbreaks appeared disconnected to Clade 3, causing new outbreaks in the Midwest and Southeastern states [19]. Phylogenetic analysis of those viruses, at the time, noted a relationship to a single Clade 2 isolate in 2016 (the last CIV sample sequenced in South Korea, SouthKorea/20170110-F1/2016) but with a sufficient branch length that makes inferences of source and relatedness difficult. These secondary outbreaks of CIV in the US now form a distinct Clade 4 (**Fig. 1**). Our newer data also reveals multiple genetically diverse virus lineages in Clade 4 that map to unique and epidemiologically disconnected outbreaks. Here we extend on these Clade 4 viruses in the US by analyzing a series of additional outbreaks occurred from mid-2017 through 2018. This includes clusters of cases due to genetically similar viruses in California, Florida, Ohio, and Kentucky, as reported previously and expanded upon here (**Fig. 1**). Another lineage involved infected dogs first seen in 2018 in California (near San Jose) and subsequently in New York (New York City). These Clade 4 viruses include several of the introductions seen in Ontario (Canada) [21]. In 2019 we identified a short-lived outbreak among imported dogs in quarantine in California which did not escape into the general dog population. Those viral sequences were not of Clade 4 but showed relationship to Clade 5 viruses observed in China (near Shanghai) and clustered with one group of the outbreak dogs in Ontario in 2018. Further into 2019 and 2020 few cases of H3N2 CIV were reported in the US and the virus appeared to have died out in North America (**Fig. 1**).

The interrelatedness of these clades, and the linear divergence structure of H3N2 CIV phylogenetics since roughly 2017 (Clades 2-4-5-6), suggest the movement of viruses between Asia and North America appears to be a hallmark of the overall epidemic.

### The epidemiology and phylogenetics of recent North American outbreaks

From the period of later in 2019 through 2020, no H3N2 CIV outbreaks were reported in North America, and there are also few reports of outbreaks in Asia. Only a single full genome H3N2 CIV sequence has been reported from China since 2019, for a virus sample collected in 2021 (with two additional HA sequences) [50]. Beginning in early 2021, an additional series of H3N2 CIV cases occurred among diverse regional outbreaks in the US, and those appeared to be more sustained and on a larger scale relative to the outbreaks occurring in the immediately preceding years. We obtained a variety of new sequences from viral infections between 2019 and 2023 by opportunistic sampling. Those covered several outbreak events, including some that appear to be ongoing at the time of preparation of this report (**Table S1)**. The sequence analysis showed the outbreaks to be widely dispersed geographically and to be temporally spread out, as some viral sequence clusters were separated by long branch lengths (**Figs. 1 and 2B**). This suggested either that we were missing key samples from our analysis, or that there were gaps in the viral transmission chains with viruses being reintroduced from outside our network.

**Figure 2.**
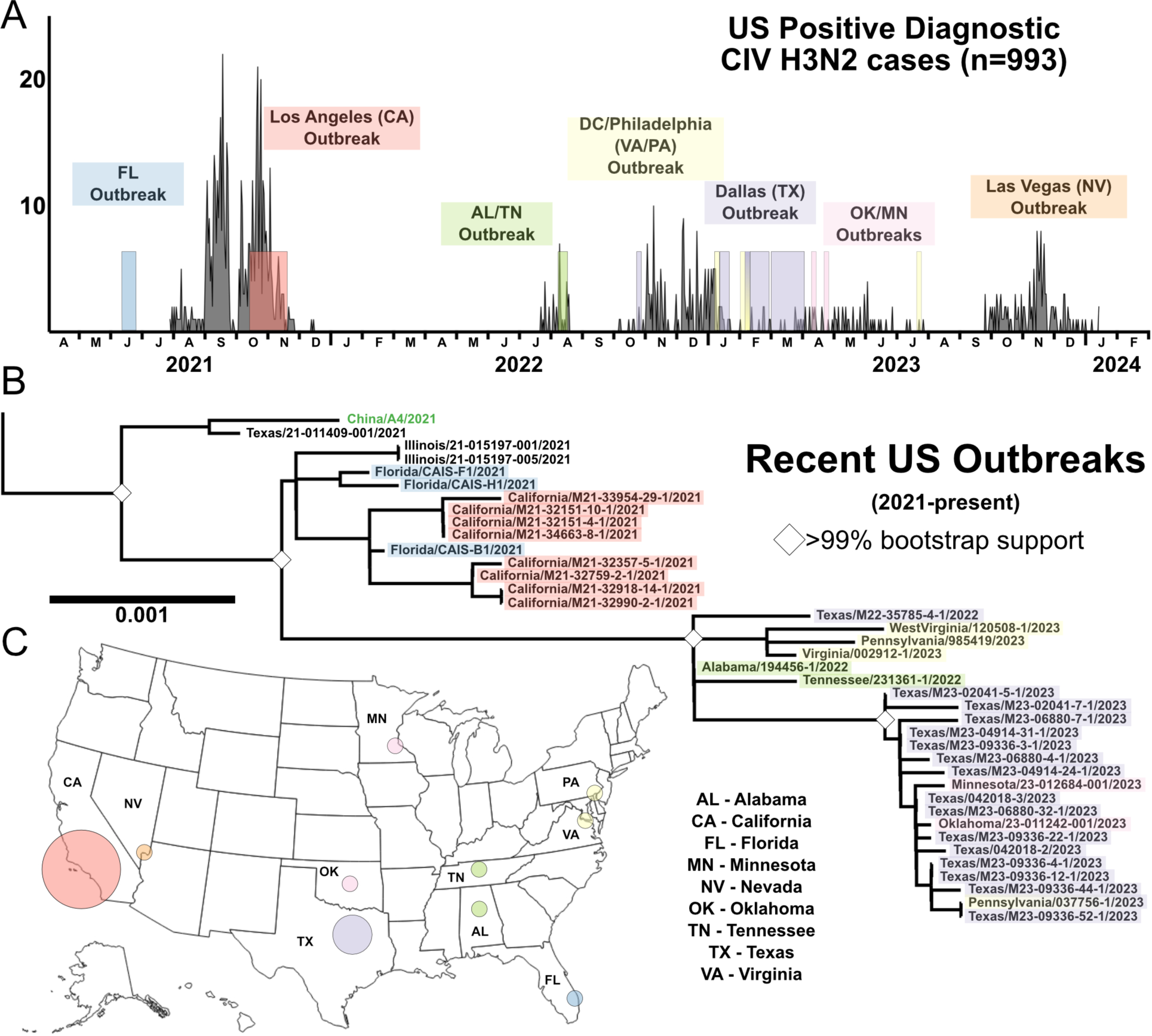
Recent circulation of H3N2 CIV in the United States, 2021-present. (A) Diagnostic positive H3N2 CIV cases (n=993). (B) Phylogenetics of full genomes among recent US outbreaks, with color highlight corresponding to timeline match with diagnostic data set and geography. (C) Geography of major US outbreaks is demonstrated with color circles corresponding to clusters in diagnostic data and phylogeny. Circles are not to scale of cases or genome count. Case data at sample geography and set to logarithmic scale available in Figure S4.

We were able to obtain US-wide diagnostic testing case data from between 2021 and early 2024, and that confirmed that our sequencing and phylogenetic analysis covered the main outbreaks that had occurred during that period (**Fig. 2A**). Gaps in our genomic data (seen as long branch lengths) mostly aligned with periods of low or absent diagnostic testing positivity, suggesting those gaps did not represent major sampling omissions. The levels of testing for the different outbreaks may differ, but there appears to be a rough concordance between the number of positive tests and the size of the outbreaks – for example, a large 2021 outbreak in Los Angeles country was represented by a large number of positive results, consistent with the County Department of Health confirmation of 1344 cases and estimates of tens of thousands of dogs being infected (case data at scale in **Fig. S4**) [51]. More recent US outbreaks in late 2023 and early 2024 centered around Las Vegas, Nevada, and we do not yet have those virus samples to compare by phylogenomic analysis. Overall, these data confirm that we have captured the major outbreaks of the virus, as well as its natural and evolutionary history since 2021 (**Fig. 2**).

We modeled the epidemiology of the virus over these major outbreaks using the different data available, including calculating estimates of the effective reproductive number (R) over time [52]. We used the diagnostic testing data set obtained to infer the scope of each local outbreak (generally within a geographical area), as well as the connections between the different outbreaks (**Fig. 3**). That analysis revealed distinct geographical outbreaks experienced fluctuations in R, which may be driven in part by localized depletion of susceptible hosts but is also expected for a host population that shows extensive contact heterogeneity - i.e. some animals being in dense and connected populations within animal shelters and kennels, while others are dispersed and disconnected as they are living in households and have many fewer contacts with other dogs. We were able to measure three predominant geographical outbreak centers: Los Angeles, California, Dallas/Fort Worth, Texas, and Las Vegas, Nevada. In each outbreak, periods of R > 1 were measured, including periods of high transmission where the R was between 2.5 and 4. Troughs of transmission with R < 1 were also observed. The largest outbreak (Los Angeles) with the greatest data volume resulted in the best resolution with mean R measures having low error variance across the timescale, where dramatic epidemiological changes to the outbreak (September 2021) can be identified. In contrast, smaller outbreaks (in Dallas and Las Vegas) with lower case data volume calculated a large degree of error and noise to make distinct temporal changes in mean R values difficult to identify with confidence. Overall, the epidemiological dynamics measured were similar to the “boom and bust” patterns seen between 2015 and 2017 in the Chicago and Atlanta areas [19].

**Figure 3.**
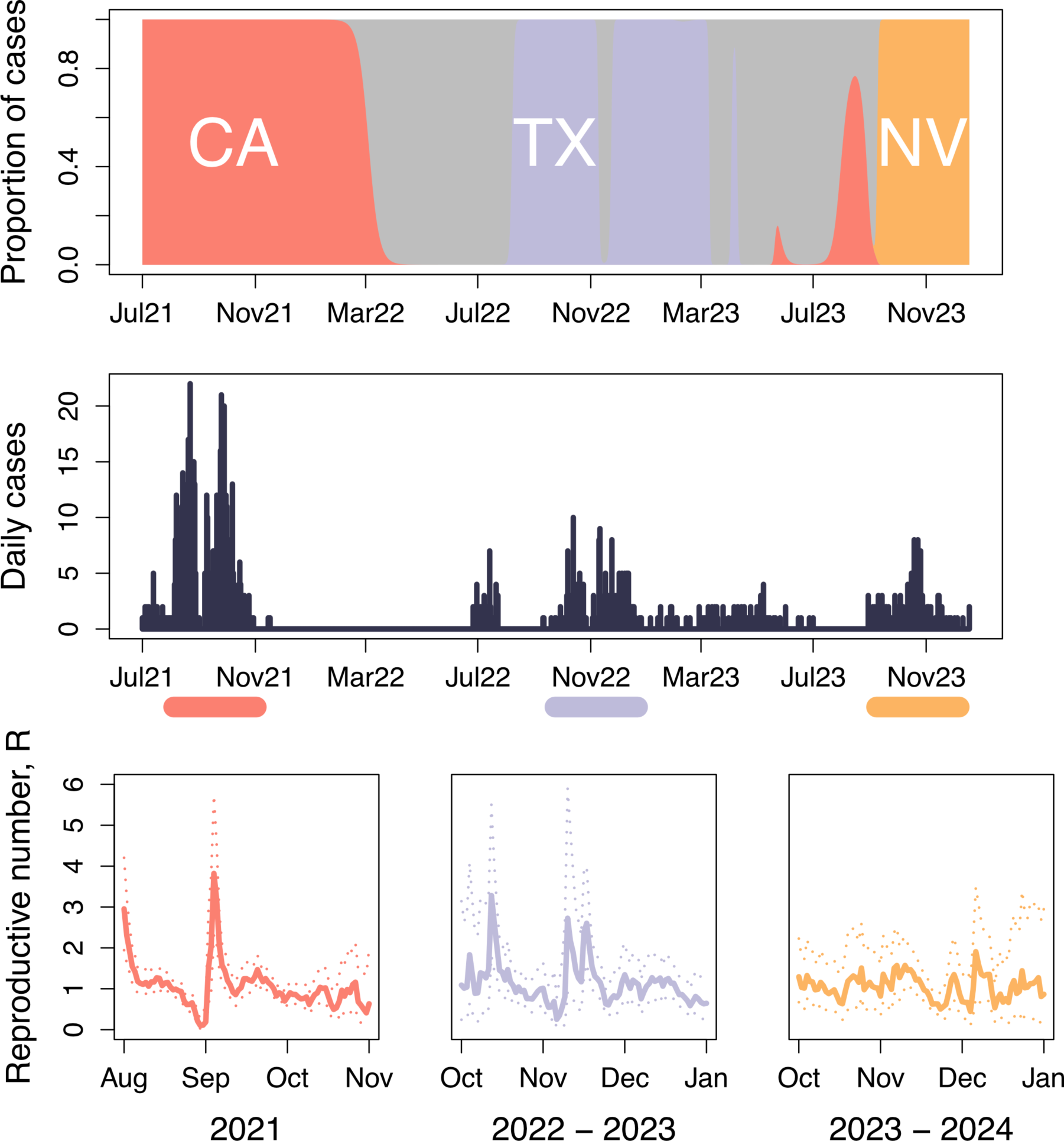
Epidemic dynamics of H3N2 CIV in the United States. Proportion of cases associated with three major outbreaks in Los Angeles (CA), Dallas/Fort Worth (TX), and Las Vegas (NV) were plotted, in addition to the number of new cases over time. Estimates of the effective reproductive number (R) were determined for each major outbreak (periods indicated by the horizontal). Solid lines show the mean estimate, while dashed lines enclose the central 95% of the posterior distribution.

### Directional transfer of H3N2 CIV between Asia and North America in recent years is result of increased virus population in Asia

A key question for understanding the epidemiology of the viruses in North America and Asia is the movement of viruses between those two regions, and in particular the directionality and frequency of any transfers that have occurred. To reveal the most likely routes of transfer, we tested a recent subset of the phylogenetic data (see Methods, Clades 4-5-6) using two models: a discrete trait diffusion model and a structured coalescent model. The implementation of discrete trait diffusion model with a Bayesian Stochastic Search Variable Selection (BSSVS) allows us to infer statistically supported rates of transition between regions across a phylogeny – i.e. showing which region which is most likely seeding infections to other regions. We also used a Structured Coalescent Model to estimate the size of virus populations in each region based on their genetic diversity, and thereby inferring the region with a larger infected population. The results of both models generally agreed, inferring similar tree topologies and transmission rates between the two regions (**Figs. 4A; S5**). Both analyses suggest five independent introductions from Asia into North America before 2021. Using the discrete trait model, we infer a higher mean rate of transitions from Asia to North America (1.4628 transitions/year, 95% HPD: [0.0332,3.6722]) than from North America to Asia (0.4968 transitions/year, 95% HPD: [6.035E-6, 1.567]), though both transition rates had strong posterior support (posterior probability of 0.99) and a Bayes Factor support (9000).

**Figure 4.**
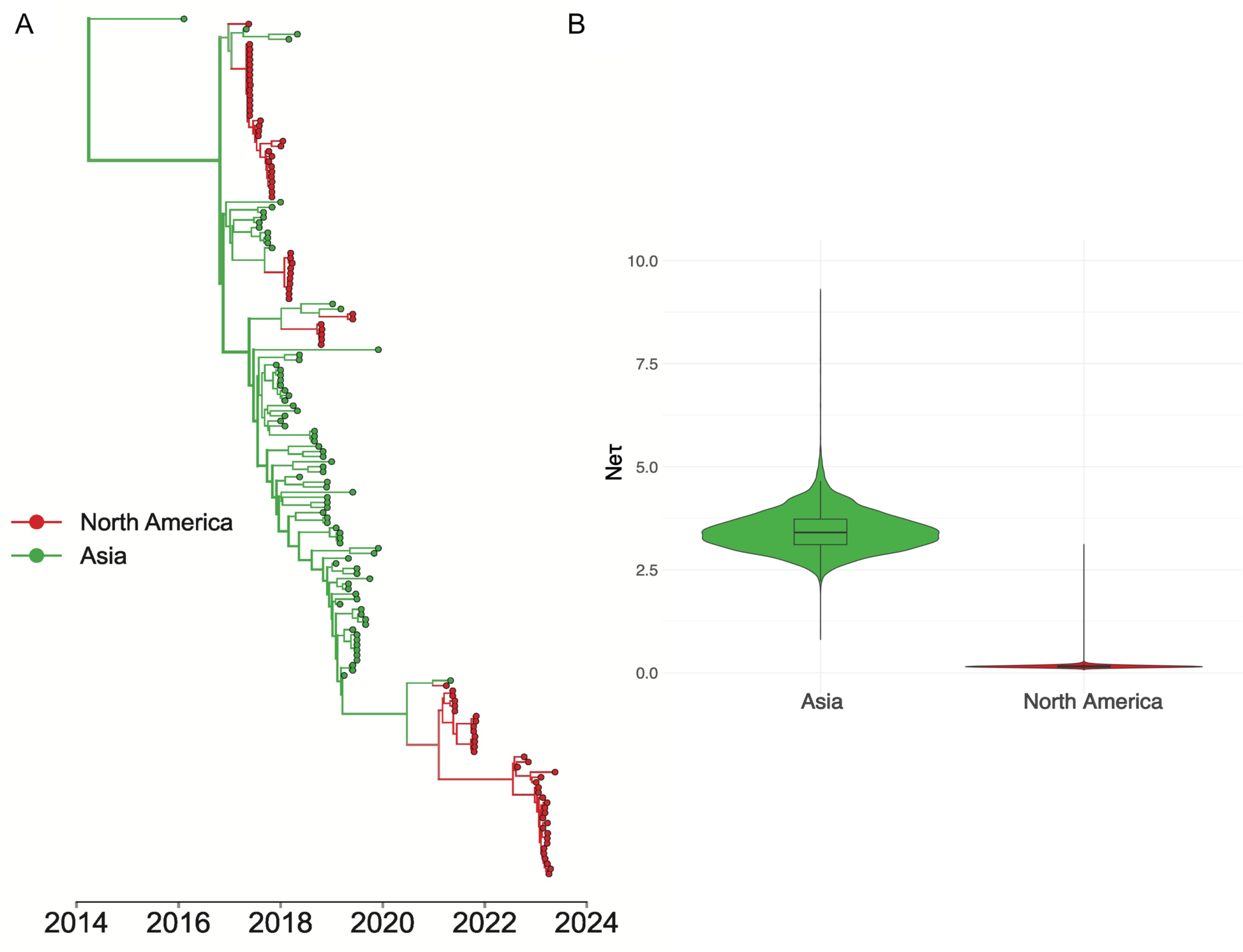
H3N2 CIV circulating in Asia act as viral sources for introductions into North America. (A) MCC tree from a structured coalescent analysis of recent subset H3N2 CIV genomes from North America and Asia. Nodes are colored by their inferred geographic region; thickness of the branches corresponds to the number of taxa which descend from the given branch. (B) Effective population size estimates for each geographic deme estimated using MultiTypeTree v8.1.0.

Although discrete trait approaches are computationally tractable, they can perform poorly in the presence of biased sampling. We therefore used a structured coalescent model to independently infer virus population sizes and rates of migrations between North America and Asia. These approaches appear to more clearly model source-sink dynamics in the presence of biased sampling [53,54], and also estimating viral effective population size (N_e_) for each region, a proxy for infected population size [55]. The mean N_e_ in Asia was 3.4438 (95% HPD: [2.6039, 4.389]), which was significantly greater than the N_e_ for North America of 0.1571 (95% HPD: [0.106, 0.2162]) (**Fig. 5B**). The higher inferred Ne matches our expectation that there is a larger virus population in Asia which acts as a source for introductions of novel lineages introduced into North America. We also estimated a backwards-in-time migration rate, which represents the rate of viruses moving to a given region A from a given region B. We estimated a higher backwards in time migration rate to North America from Asia (1.260 migrations/year, 95% HPD: [0.4474, 2.1399]) compared to the rate to Asia from North America (0.0894 migrations/year, 95% HPD [2.9079E-3, 0.2134]), consistent with the results of the discrete trait diffusion.

**Figure 5.**
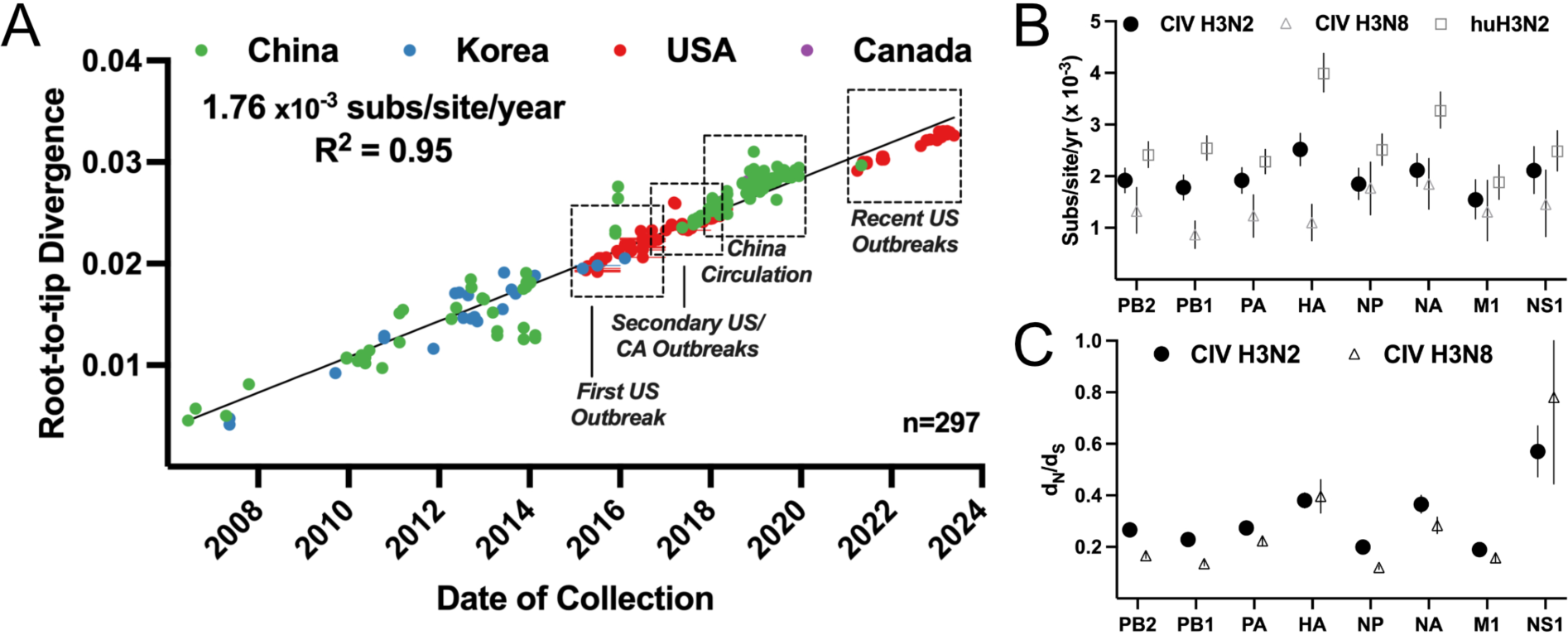
Temporal evolution of H3N2 CIV during continuous dog-to-dog circulation. (A) A root-to-tip analysis of H3N2 CIV full genomes, showing the divergence since the first common ancestor of the virus represented by the basal node of the phylogeny. This shows a consistent evolutionary rate of 1.76 x 10^-3^ substitutions/site/year. (B) Individual segment ORF substitution rates were calculated in BEAST and compared to H3N8 CIV and human seasonal H3N2. (C) Mean segment ORF d_N_/d_S_ ratios were calculated using SLAC and compared to H3N8 CIV.

### The rate of H3N2 CIV sequence evolution is consistent over host geography and time

To examine the temporal nature of the full genome data set, we used well dated samples (YYYY-MM-DD) to analyze the maximum likelihood genome phylogeny, plotting the root-to-tip length data against sampling date (using the TempEst algorithm). That showed a consistent clock-like rate that averaged 1.76 x 10^-3^ substitutions/site/year (R^2^ = 0.95) over all outbreaks across two continents, and since the emergence of H3N2 CIV in dogs around 2004 (**Fig. 5A**). That consistent clock-like evolution revealed no evolutionary behaviors that might indicate pronounced selections (e.g., bursts of adaptive evolution) on the virus in either its apparent reservoir among dogs in Asia or during outbreak spread within North America. Attempts to analyze the clock by distinct geography or timescale windows showed no noteworthy variation from the full data set mean (data not shown). Any further variance in individual segments was also not seen (**Fig. S3**).

The nucleotide substitution rate for each genome segment was further determined using a Bayesian coalescent method in BEAST. Among the H3N2 CIV segment ORFs, substitution rates were largely similar – with a higher rate for the HA gene segment (rising to significance against 5 of 7 other segments, save NA and NS1) (**Fig. 5B**). These rates are largely similar to those seen for another influenza virus previously seen in dogs (H3N8 equine-origin CIV, analysis from Wasik *et al*. [12]), though higher rates were seen in H3N2 CIV among the PB1, PA and HA segments (1.782 vs 0.87, 1.917 vs 1.237, and 2.519 vs 1.102, all 10^-3^ substitutions/site/year, respectively). The rates seen in both the H3N2 and H3N8 CIVs were lower in nearly all segments than those observed in seasonal H3N2 human influenza virus, with greatest pronouncement and significance relative to the HA and NA glycoproteins [56]. To generate a rough proxy of natural selection in the H3N2 CIV during the 20 years of dog-to-dog transmission, we used the model statistical method SLAC (Single-Likelihood Ancestor Counting) [57] to determine the mean d_N_/d_S_ ratio for each segment (**Fig. 5C**). The greatest potential signals of positive selection were seen in the HA, NA, and NS segment ORFs. The d_N_/d_S_ ratios observed in CIV were generally higher than previous measures of H3N8 CIV (2003-2016) evolution, apart from HA and NS.

### Cumulative fixation of non-synonymous mutations during H3N2 CIV evolution

To understand the likely functional effects of the viral evolution, we identified non-synonymous mutations that became fixed during H3N2 CIV evolution (**Figs. 1 and 6**). Most clustered at key nodes within the phylogenetic tree, appearing following likely bottlenecks during transfers to new geographical regions. The overall evolution of the virus is re-reviewed in **Fig. 6**, and key non-synonymous mutations that became fixed during the evolutionary pathway are highlighted. For example, 13 coding mutations became fixed within 7 of the 8 gene segments of the virus during the emergence of Clade 2 in South Korea, and additional groups of non-synonymous mutations (ranging in numbers from 3 to 15 over the entire genome) became fixed during the subsequent clade-defining events (**Fig. 6, Tables S3-4**).

**Figure 6.**
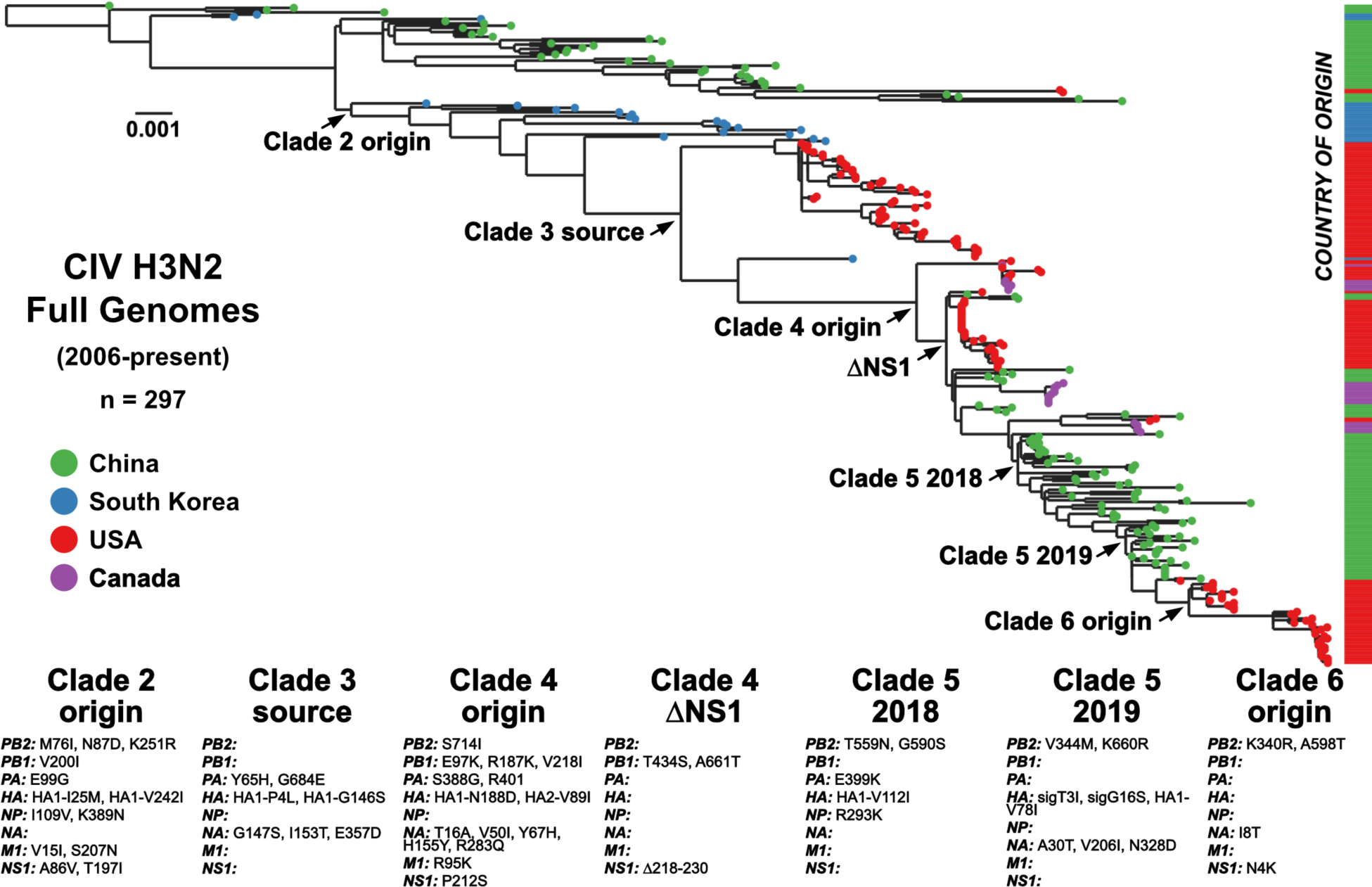
Fixed mutations at key transitional nodes and international transfer events. (A) A ML tree of H3N2 CIV genomes with geographic sampling sources coded on tips (green = China, blue = Korea, red = US, purple = Canada) and on corresponding bar to right of tree. (B) Noted start of Clade 2, circulating in South Korea. (C) Transition of Clade 2 to Clade 3, international introduction of US outbreaks, 2015-2017. (D) Start of Clade 4, after last known South Korean isolate. (E) A major early genetic bottleneck of Clade 4 by NS1 truncation. (F) Early convergence in Clade 5, ∼2018 circulation in China. (G) Later convergence in Clade 5, ∼2019 circulation in China. (H) Start of Clade 6, with US outbreaks in 2021-present.

Mutations span the genome and appear in multiple functional domains of the influenza gene products. A notable fixation was the truncation of the NS1 protein at the C-terminus during Clade 4 evolution. A truncation of NS1 has previously been observed in the H3N8 subtype in horses and dogs and was related to host-specific adaptation of innate immunity [58,59]. Several changes in PB2 (S714I) and PA (E327K and S388G) have previously been identified during mammalian adaptation of the polymerase subunits, related to ANP32A co-factor utilization [23,60].

### HA-specific mutations

The HA protein controls receptor binding, cell membrane fusion, and is the major antigenic determinant of influenza, so that mutations in the HA may play key roles in controlling CIV biology and epidemiology. We identified the major fixed HA mutations of H3N2 CIV following the establishment of Clade 4 (**Fig. 7A, Table S3**) and mapped their position onto the HA monomer structure (**Fig. 7B**). Following the last reported South Korean-origin genome, several nonsynonymous mutations have fixed in the HA gene of CIV lineages and clades. First, upon the establishment of Clade 4, there fixed an asparagine to aspartic acid mutation (N188D, in H3 numbering) at the top of the HA1 head near the sialic acid receptor binding site (RBS). This mutation was discussed in Chen *et al* as potentially playing a role in clade-specific phenotypes [49]. That Clade 4 transition also fixed a stalk mutation (HA2-V89I). During Clade 5 evolution during circulation in China, two major nodes were identified in phylogenetic analysis. Upon circulation in 2018, there fixed another HA1 mutation from valine to isoleucine (V112I). Later in 2019, there fixed both an HA1 mutation (V78I) as well as a glycine to serine change in the final residue of the signal peptide (G*sig16*S). Clade 6 represents all US outbreaks after 2021 where lineages likely resulted from multiple independent introductions, therefore unique HA fixed mutations can be seen with temporal and geographic patterns (**Table S4**). An HA2 mutation was seen fixed in the earliest US samples in 2021 (R82K), while an HA1 asparagine to threonine change was seen starting in 2022 viruses (N171T, in the RBD). In the most recent Clade 6 viruses in the US (2023), we observe regional subclade lineages with unique HA fixations (**Fig. 7**). In the Mid-Atlantic region (around Washington DC and Philadelphia, Pennsylvania) viruses have an HA2 mutation (S113L) in addition to an isoleucine to methionine (I245M) change in the HA1 head near the RBD. In contrast, viruses found in the subclade around Dallas, Texas, contain a HA1 mutation located in a region that contains antigenic site D in human H3 viruses (I245M).

**Figure 7.**
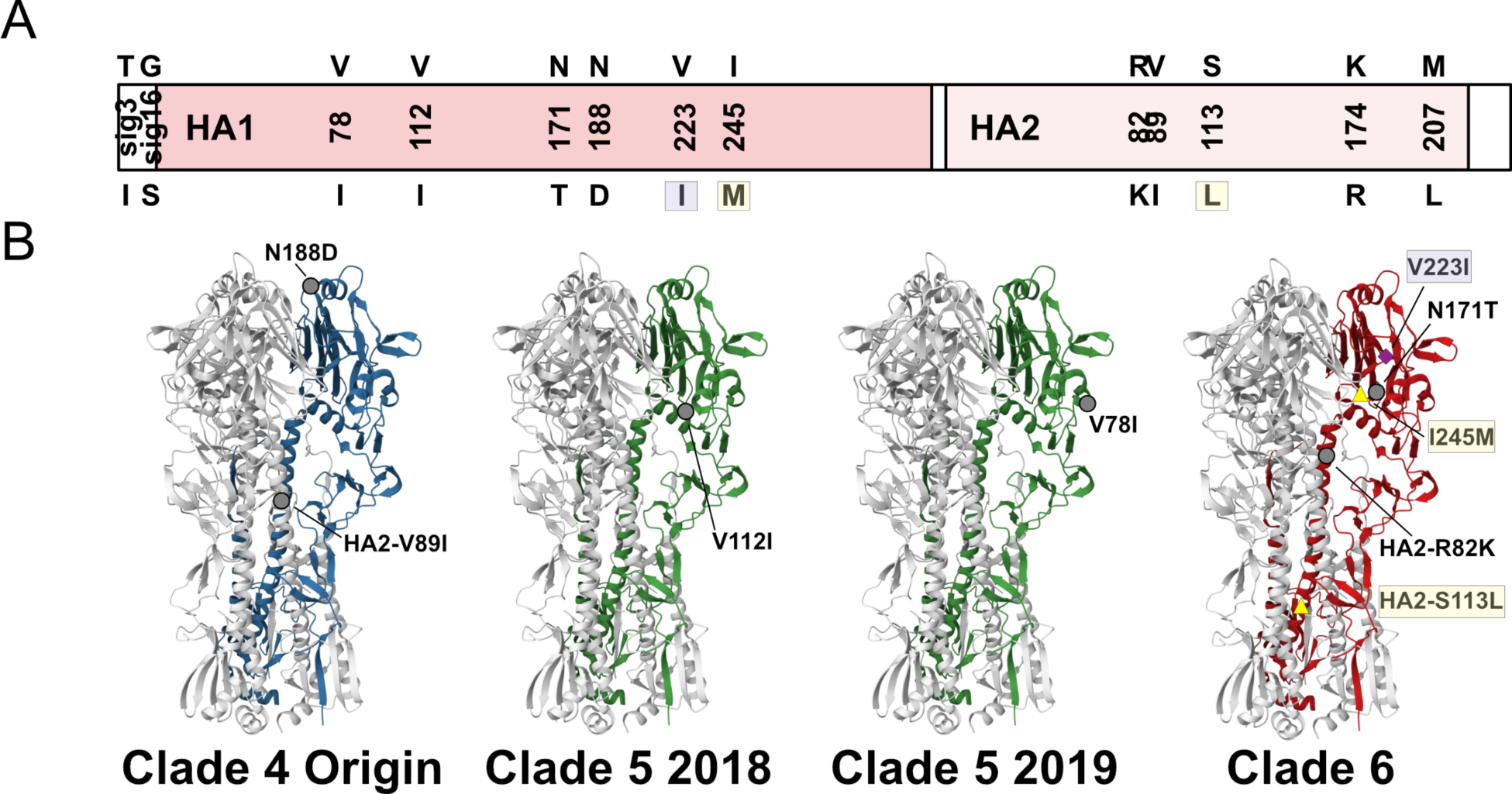
Recent molecular evolution of H3N2 CIV hemagglutinin. (A) Schematic of the HA ORF showing with nonsynonymous mutations fixed since the last Clade 2 isolate, SouthKorea/20170110-1F1/2016. HA1 and HA2 positions follow H3 numbering. Signal peptide sequence residues are noted as sigX. (B) The location of key HA1 and HA2 mutations on the monomer of HA, occurring during key transitions in the recent evolution of H3N2 CIV. Mutations specific to most recent Clade 6 US subclades (Mid-Atlantic, yellow and Texas, purple) are highlighted.

## Discussion

Here we further explain how the H3N2 canine influenza virus (CIV) emerged, evolved, and spread around the world during the 20 years that it has been circulating in dogs, providing an example of an influenza virus switching hosts to cause a sustained epidemic in a mammalian host population. Our results emphasize the interconnections between viruses circulating in different regions of Asia and those in North America. We show that around 2017 the H3N2 CIV was reduced to a single lineage that has since expanded to include viruses in mainland China, the United States and Canada. The data also strongly suggests that there were two introductions into the USA in 2021 and in 2022. The first lead to outbreaks during 2021 that started in Florida and then caused a large outbreak in the Los Angeles, California area that continued for several months, but which died out by the end of that year. The second introduction occurred around June or July of 2022 that resulted in a sustained outbreak Dallas/Fort Worth, Texas, which then spread to cause a series of outbreaks in many areas of the USA, which have continued to the present (**Fig. 2**).

### Overall and recent epidemiology

The results again confirm that the epidemiology of this virus in dogs in North America is outbreak-driven and constrained by the host population structure. Those are characterized by rapid transmission within geographic areas where the virus spreads among group-housed dogs in kennels, animal shelters, and day-care settings. The largest and longest duration outbreaks have occurred within larger metropolitan areas, where circulation has been sustained for up to several months, although all observed outbreaks ultimately died out after between 1 and 6 months (**Fig. 2**). The more recent epidemiology in North America was revealed by the analysis of nation-wide diagnostic testing data, which showed the duration, intensity, and locations of the major outbreaks that occurred (**Figs. 2 and 3**). That analysis was also compared to parallel phylogenetic and phylogeographic studies based on full genome sequencing data. Although both sets of data resulted from opportunistic sampling, the patterns revealed were similar, indicating that they would explain the virus spread and evolution in North America during this period.

Little recent data is available about the viruses in Asia and their circulation patterns, with only a single full H3N2 CIV sequence being available since 2019, so it is not possible to compare the details of the recent epidemiology or evolution of the viruses in Asia and North America. However, it appears that during the last several years the viral population has been distributed between China and North America (with no virus being reported from South Korea since around 2017). Our analysis suggests a significantly greater likelihood of transmission from Asia to North America than the reverse (**Fig. 4**).

The effective reproduction number (R) for the H3N2 CIV was estimated from the data, and varied widely depending on the structure of the dog population. Within the large metropolitan centers, the R was between 1 and 4, as also revealed by the rapid and sustained spread of the virus within more densely housed dogs. However, ultimately within each population the recovered animals would become resistant due to the development of protective immunity, so without the addition of large numbers of new susceptible animals the Susceptible Infected Recovered (SIR) dynamic would apply, and the virus would die out naturally.

A common prediction is that transmission of a virus in a new host will select for host-adaptive mutations which would result in more efficient infection, replication, and transmission – often referred to as “gain-of-function” mutations. While the H3N2 CIV has acquired many changes with the potential to be host-adaptive during its thousands of exclusively dog-to-dog infections and transmissions, but we do not yet see clear evidence of CIV being able to spread more effectively in dogs within dense populations, or gaining an increased ability to transfer between geographically separate population centers.

### H3N2 CIV evolution and possible host adaptation

The H3N2 CIV has likely undergone over 1,000 exclusively dog-to-dog infections and transfers since it emerged, and there has been a linear acquisition of genetic change since 2005. While some adaptive changes likely arose during the first series of transfers of the H3N2 CIVs in dogs, those are difficult to define as the sequence of the directly ancestral avian virus is not known [16]. While several mutations in H3N2 CIV are in viral genes and proteins associated with host adaptation in other influenza spillovers (**Tables S3-4**), we do not have clear evidence that those have resulted in gain of function for the virus [49,61]. In other studies experimental passage of as few as six ferret-to-ferret transfers were sufficient to make high pathogenicity H5N1 influenza viruses more transmissible in that new host [62,63], while for SARS CoV-2 several more transmissible variants have arisen during the first years of spread in humans [64–67]. A possible explanation is that the complex environmental pressures of the canine population may be constraining the viral evolution, since most spread and infections have occurred in dense populations of animals in close contact, but which result in SIR-mediated die outs. The mutations selected would therefore differ from those that would favor the long-distance transfers between population centers required to sustain long-term transmission. Properties that may favor the second type of spread might include persistent infections, prolonged shedding, virus stability and fomite-mediated transfer, as suggested by life-history tradeoff evolutionary theory [68].

### Are H3N2 CIVs a risk to humans and might those risks be changing?

The risk of emergence of the H3N2 CIV into additional mammalian hosts (including humans) is a concern, but currently the threat is unknown. While swine appear to be intermediate host that allows influenza viruses to jump onward into humans to cause sustained epidemics on at least a few occasions, other mammalian-adapted influenza viruses (in horses, seals, cows, mink) have so-far not proved to be clear threats to humans [3]. The changes in the H3N2 CIV have included previously identified mammalian-adaptive sites [23], some experimentally confirmed as having biological effects [49], and the H3N2 CIV appears to lost the host range for at least some avian hosts [69]. Other concerning adaptive changes that underlie known human adaptation and pandemic risk (including NP changes to evade MxA/BTN3A3 restriction factors, and HA RBD changes to the 226/228 dyad) are absent, as the H3N2 CIV remains ‘avian-like’ [70–72].

Changes have occurred in NA domains related to ligand binding, but the constellation of changes do not include any known inhibitor-resistance markers, suggesting antiviral use in zoonotic exposure would be efficacious [73]. Cross-species transmission of H3N2 CIV into cats has occurred in Korea and US [74,75]. In a serosurvey of race horses in China, animals in several in riding clubs were positive for H3N2 CIV, with infection being strongly associated with exposure to dogs [76]. To date no natural human infections with either H3N8 or H3N2 CIV have been reported. Following the 2015 US outbreak of H3N2 CIV, a human spillover risk assessment was performed on a Clade 3 isolate, examining for receptor utilization, replication kinetics in human cells, and antibody cross-reactivity, and concluded that H3N2 CIV posed a low risk for human populations [61].

Besides point mutations, reassortments of influenza viruses allow mixing of viruses to find favorable genetic combinations [77,78]. No reassortant influenza viruses were seen in dogs in the United States (**Figs. S1-2**). H3N2 CIV reassortants with human, swine, and avian influenzas have been reported in Korea and China [79–82], but none have generated sustained transmission chain lineages and the fitness effects for either dogs or humans are unknown.

While the H3N8 and H3N2 CIVs both circulated in dogs in the USA during 2015 and 2016, the two strains were not found within the same population so no recombinant opportunities likely occurred [12,19]. The lack of reassortant viruses in North American dogs suggests differences in host population ecology or epidemiology of the CIV and the other viruses leading to fewer mixed infections. It is unknown whether additional viral strains including high pathogenicity avian influenza (HPAIV) H5N1 clade 2.3.4.4b may create reassortant events, but additional screening of domestic animals for influenza viruses is likely warranted [83–86].

### Summary and Conclusions

Respiratory viruses emerging to cause human pandemics readily spread through host populations, crossing oceans within days and overwhelming control measures [34,87,88]. Here we show that the spread of emerging epidemic diseases in animal populations may differ significantly from the patterns seen in humans, even though the transmission potential for each virus is similar. The different outcomes are mostly due the different population structures and inter-connectedness of different wild or domesticated animals [13]. There are around 900 million domestic dogs world-wide, and many live in very close proximity to their human owners, presenting a special opportunity for high levels of human exposure [43]. While the H3N2 CIV has circulated widely in Asia and North America, and undergone several transfers between those regions, it has not been reported from Europe, Australia, Africa or other regions with large dog populations and veterinary disease testing. The observations that the H3N2 CIV has repeatedly died out in North America, and that the H3N8 CIV also died out in North American dogs in 2017 with little directed intervention [12,19], indicates that modest interventions such as animal symptom screening, quarantine, and slowing the movement of infected dogs would allow the H3N2 CIV to be both eradicated from the dog population, while modest levels of testing or quarantine of imported dogs would prevent the virus from being re-introduced. We propose that the epidemiological and evolutionary patterns seen here may prove a useful comparison with viruses in other hosts – horses, swine, seals, cows – where interventions at key points could also result in them being eradicated with relatively little effort.

## Methods and Materials

### H3N2 Sample Collection

Virus samples were obtained from various diagnostic centers following routine passive diagnostic testing for respiratory disease in dogs (listed in Table S1, red). Samples were received as either nasal or pharyngeal swab material or extracted total nucleic acid (tNA) first utilized for quantitative reverse transcription-PCR (qRT-PCR) positivity. Some additional samples were first isolated in embryonated chicken eggs or MDCK cells for virus isolation. vRNA was extracted from clinical swab samples and virus isolation using the QIAamp Viral RNA Mini kit. Purified vRNA or tNA was then either directly used for influenza multi-segment RT-PCR or stored at -80°C.

### Generation of Influenza Viral Sequences

Virus genomes from samples were generated as cDNAs using a whole genome multi-segment RT-PCR protocol, described previously [12,89]. A common set of primers (5′ to 3′, uni12a, GTTACGCGCCAGCAAAAGCAGG; uni12b, GTTACGCGCCAGCGAAAGCAGG; uni13, GTTACGCGCCAGTAGAAACAAGG) that recognize the terminal sequences of the influenza A segments were used in a single reaction with SuperScript III OneStep RT-PCR with Platinum *Taq* DNA polymerase (Invitrogen). Following confirmation by gel electrophoresis, viral cDNA was purified either by standard PCR reaction desalting columns or with a 0.45X volume of AMPure XP beads (Beckman Coulter). Libraries were generated either by Nextera XT with 1 ng of cDNA material, or Nextera FLEX with 150-200 ng (Invitrogen). Libraries were multiplexed, pooled, sequenced using Illumina MiSeq 2 X 250 sequencing. As most samples were isolated from direct nasal swabs and may contain intra-host viral diversity at sequencing depth, raw reads were deposited in the Sequence Read Archive (SRA) at NCBI.

Consensus sequence editing was performed using Geneious Prime. Paired reads were trimmed using BBduk script (https://jgi.doe.gov/data-and-tools/bbtools/bb-tools-user-guide/bbduk-guide/) and merged. Each sequence was assembled by mapping to a reference sequence of a previously annotated H3N2 isolate (A/canine/Illinois/41915/2015(H3N2)). Consensus positions had read depth >300 and >75% identity.

### Diagnostic Testing Data and R Estimation

Diagnostic qRT-PCR tests positive for H3N2 CIV were obtained, deidentified save test date and zip code, from a commercial vendor. The total data set (n=993) spans from 2021.07.27 to 2024.01.12, and identifies cases in 17 US states by zip code. R estimation from daily case data followed the standard methodology described in [52]. The generation time was assumed to follow a discretized version of a gamma distribution with a mean of 3.5 days and a variance of 2 days. The temporal window over which R was assumed constant was 7 days.

### Phylogenetic Analysis

H3N2 CIV nucleotide sequences were downloaded from the NCBI Influenza Virus Database and were compiled with generated consensus genomes for this study and organized in Geneious Prime. We examined the larger database collection of H3N2 present in dog hosts for both inter-and intra-subtype reassortants using RDP4 (seven methods: RDP, GENECONV, Bootscan, MaxChi, Chimaera, SiScan, 3seq) and excluded those found to be a statistically significant outlier in two or more methods [90]. The data set with full genome coverage (n=297) is provided in Table S1. Sequences were manually trimmed to their major open reading frames (PB2: 2280nt, PB1: 2274nt, PA: 2151nt, HA: 1701nt, NP: 1497nt, NA: 1410nt, M1: 759nt, and NS1: 654-693nt) and either analyzed separately or concatenated with all other genome segments from the same virus sample. Nucleotide sequences were aligned by MUSCLE [91] in the Geneious Prime platform. Maximum likelihood (ML) phylogenetic analysis was performed by either PhyML or IQ Tree [92,93], employing a general time-reversable (GTR) substitution model, gamma-distributed (Γ4) rate variation among sites, and bootstrap resampling (1000 replications). ML trees were visualized and annotated using FigTree v1.4.4 (tree.bio.ed.ac.uk/software/figtree/). An additional reassortment analysis for all 297 aligned H3N2 CIV sequences was performed using Treesort (github.com/flu-crew/TreeSort). Reassortment events were inferred on each branch of the phylogeny using HA as the fixed reference tree.

Temporal signal was assessed by a regression root-to-tip genetic distance against date of sampling using our ML tree and the TempEst v.1.5.3 software [94]. Accurate collection dating to the day (YYYY-MM-DD) was utilized for all samples where this information was available. Sampling dates used for all isolates are listed in Tables S1.

### Bayesian Phylodynamic Analysis

We used a Bayesian coalescent approach to better estimate our phylogenetic relationships, divergence times, and population dynamics. Analyses were performed in BEAST v.1.10.4 [95], where Markov chain Monte Carlo (MCMC) sampling was performed with a strict clock, a general time-reversable (GTR) substitution rate with gamma distribution in four categories (Γ4), set for temporal normalization by sample collection date (YYYY-MM-DD to best accuracy), and assuming a Bayesian Skyline Plot (BSP) prior demographic model. Analyses were performed for a minimum of 100M events, with replicates combined in Log Combiner v1.10.4. Outputs were examined for statistical convergence in Tracer v1.7.2 (effective sample size [ESS] ≥200, consistent traces, removed burn-in at 10-15%) [96].

For the discrete trait diffusion model [97] and the structured coalescent model [98], we used a subset of diverged lineage genomes (n = 169) starting with the last Korean isolate (SouthKorea/20170110-1F1/2016) in Clade 2 and related lineages in Clades 4, 5, and 6. This data set was annotated for two geographic locations (Asia or North America) based on sample collection. These annotations for geographic region were used as discrete traits for discrete trait diffusion modelling and as demes for structured coalescent modelling. We first performed a discrete trait diffusion model for the sequences using BEAST v1.10.4. We used an HKY nucleotide substitution model with gamma distributed rate variation among sites, a lognormal relaxed clock model, and GMRF skyride model [99–101]. We ran six independent MCMC chains of 50 million steps, logging every 5,000 steps. Outputs were examined for statistical convergence in Tracer v1.7.2 (effective sample size [ESS] ≥200, consistent traces, removed burn-in at 10%). The three runs with the highest ESS values were selected and combined using LogCombiner v1.10.4. The last 500 posterior sampled trees from the combined runs were used for as an empirical tree set to preform discrete trait diffusion analysis. We performed an ancestral state reconstruction to determine the discrete trait states across all branches of the phylogeny. We used the Bayesian stochastic search variable selection method (BSSVS) to estimate the most parsimonious asymmetric rate matrix between discrete traits [97]. We used a Poisson mean prior of 1 non-zero rate. The Bayes factor support for the rates were calculated using SpreaD3 v0.9.7.1 [102]. The maximum clade credibility (MCC) tree was summarized with the program TreeAnnotator v1.10.4 using the combined tree files for the runs with highest ESS value using a posterior sample of 10,000 trees. The structured coalescent analysis was implemented using the MultiTypeTree v 8.1.0 package available in BEAST v2.7.6 [103]. These models estimate an effective population size (N_e_) in each deme, and asymmetric migration rates between demes [98]. We calculate the asymmetric migration rates as “backwards-in-time” rates which parameterize the rate of movement of ancestral tree lineages between demes backwards in time. This represents the probabilities per unit time of individual members of a particular deme having just transitioned from some other deme. We also calculate the “forwards-in-time” rates which parameterize the (constant) probability per unit time that an individual in some deme will transition to a new deme. The relationship between the backwards and forwards migration rates is given by:

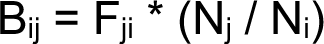

where B_ij_ is the backwards-in-time rate from deme i to deme j, F_ij_ is the forwards-in-time rate from deme i to deme j, and where N_i_ and N_j_ are the effective population sizes of these two demes [54]. We used an asymmetric model of migration with a lognormal prior distribution for the rate with a mean of 1 and upper limit of 20 as well as a uniform prior distribution for the Log Population size tau (Effective population size) between 0.001 and 10000. We ran six independent MCMC chains of 100 million steps, logging every 10,000 steps. Run convergence was assessed using Tracer 1.7.2 (effective sample size [ESS] ≥200, consistent traces, removed burn-in 10%). The three runs with the highest ESS values across parameters were combined using LogCombiner v1.10.4. The MCC tree was summarized for the tree files of the three runs with the highest ESS using the program TreeAnnotator v1.10.4 for a posterior sample of 10,000 trees. Visualization for MCC trees was achieved using the BALTIC python package (https://github.com/evogytis/baltic). Branches of the phylogenies were colored based on annotations for the most probable ancestral state or deme for the MCC tree of the discrete trait diffusion model analysis and structured coalescent analysis respectively.

### Analysis of Selection Pressures

The relative numbers of synonymous (d_S_) and nonsynonymous (d_N_) nucleotide substitutions per site in each ORF of each segment were analyzed for the signature of positive selection (i.e. adaptive evolution, p<0.05 or >0.95 posterior) using SLAC (**S**ingle-**L**ikelihood **A**ncestor **C**ounting) within the Datamonkey package datamonkey.org, [57]).

### Hemagglutinin Structural Modeling

HA structures and positions of CIV mutations were visualized in Mol* from the RCSB Protein Database (PDB). The structure of A/HongKong/1/1968 (4FNK) was used as an H3 model.

### Graphing and Statistical Analysis

Graphs in Figures 2 and 5 were generated in GraphPad Prism v.10. Figure 3 which was generated by R version 4.2.3 without the use of any additional packages.

## Data Availability

All generated full genome sequence data have been submitted to NCBI under BioProject PRJNA971216. Raw sequence reads from amplified direct swab samples were deposited in the Sequence Read Archive (SRA). Consensus genome sequences were submitted to Genbank, with accession numbers listed in Table S1, in addition to public sequences retrieved and employed. All BioSample and SRA accession numbers of NGS of direct nasal swabs generated in this study are listed in Table S2.

## Acknowledgements

We wish to thank multiple Veterinarian sources for narratives of observed outbreaks, in addition to the collection and sharing of diagnostic samples. We thank the University of Wisconsin Veterinary Diagnostic Lab (WVDL) for support in diagnostic testing for H3N2 CIV. We thank the Cornell Animal Health Diagnostic Center (AHDC), and particularly Diego Diel and Lina Covaleda, for support in diagnostic testing for H3N2 CIV and virus isolation. We thank Joseph Flint, Guillaume Claud Reboul, and Kelly Sams of the Goodman Lab for technical support in library preparation and sequencing. Significant lab support was provided by Wendy Weichert.

We thank Timothy Vaughan, ETH Zurich, for helpful discussions on structured coalescent models. We thank Eddie Holmes, University of Sydney, for key feedback and insight during drafting of this manuscript. CML is currently an employee of Antech Diagnostics, Mars Petcare Science and Diagnostics, however this organization did not participate in funding this work. This work was partially funded by the Influenza Division of the Centers for Disease Control (CDC), Animal-Human Interface program, under contract #75D30121P12812 to CRP and LBG. The content and commentary in this manuscript is solely the responsibility of the authors and does not necessarily represent the official views of the CDC.

