## Supplemental Tables/Figures for "The Evolution and Epidemiology of H3N2 Canine Influenza Virus After 20 Years in Dogs"

**SUPPLEMENTAL TABLES AND FIGURES – WASIK ET AL.**

| **Segment** | **Position** | **Clade 1** | **Clade 2** | **Clade 3** | **Clade 4** | **Clade 5** | **Clade 6** | **Notes/**  **References** |
| --- | --- | --- | --- | --- | --- | --- | --- | --- |
| PB2 | 76 | M, I | ***I*** | I | I | I | I |  |
| PB2 | 87 | N, D | ***D*** | D | D | D | D |  |
| PB2 | 175 | R | R | R | R | R, ***K*** | K |  |
| PB2 | 251 | K | ***R*** | R | R | R | R |  |
| PB2 | 340 | K | K | K | K | K | ***R*** |  |
| PB2 | 344 | V | V | V | V | V, ***M*** | M | Pol oligomer |
| PB2 | 559 | T | T | T | T, N | ***N*** | N | T-cell epitope[1] |
| PB2 | 590 | G | G | G, S | G, S | ***S*** | S | Ligand binding |
| PB2 | 598 | A | A | A | A | A | ***T*** | Virulence (H7N9)[2] |
| PB2 | 660 | K | K | K | K | K, ***R*** | R | Pol oligomer[3] |
| PB2 | 714 | S | S | S | ***I*** | I | I | Virulence/Host-range, Pol oligomer[4–6] |
| PB1 | 11 | K | K | K | K | K, ***R*** | R | Pol oligomer[7] |
| PB1 | 97 | E | E | E | ***K*** | K | K |  |
| PB1 | 187 | R | R | R | ***K*** | K | K |  |
| PB1 | 200 | V | ***I*** | I | I | I | I |  |
| PB1 | 216 | S | S, ***N*** | N | N | N | N |  |
| PB1 | 218 | V | V | V | ***I*** | I | I |  |
| PB1 | 434 | T | T | T | ***S*** | S | S |  |
| PB1 | 661 | A | A | A | ***T*** | T | T |  |
| PA | 65 | Y | Y, ***H*** | H, Q | H | H | H | Pol oligomer[8] |
| PA | 99 | E | ***G*** | G | G | G, ***E*** | E | Pol oligomer |
| PA | 241 | C | C, ***Y*** | Y | Y | Y | Y |  |
| PA | 327 | E | E, ***K*** | K | K | K | K | Pol oligomer[4] |
| PA | 388 | S | S | S | ***G*** | G | G | Pol oligomer[4] |
| PA | 399 | E, K | E | E | E, K | ***K*** | K |  |
| PA | 401 | R | R | R | ***K*** | K | K |  |
| PA | 684 | G | G, ***E*** | E | E | E | E |  |
| HA | Sig3 | T | T | T | T | T, ***I*** | I |  |
| HA | Sig14 | G | G | G | G | G, ***S*** | S |  |
| HA1 | 4 | P | P, ***L*** | L | L | L | L |  |
| HA1 | 25 | I | ***M*** | M | M | M | M |  |
| HA1 | 78 | V | V | V | V | V, ***I*** | I |  |
| HA1 | 112 | V | V | V | V, I | ***I*** | I |  |
| HA1 | 146 | G | G, ***S*** | S | S | S | S |  |
| HA1 | 188 | N | N | N | ***D*** | D | D |  |
| HA1 | 242 | V | ***I*** | I | I | I | I |  |
| HA2 | 82 | R, K | ***R*** | R | R | R, ***K*** | K |  |
| HA2 | 89 | V | V | V | ***I*** | I | I |  |
| HA2 | 207 | M | M | M | M | M, ***L*** | L |  |
| NP | 109 | I | ***V*** | V | V | V | V | Host-range (H1N1pdm)[9] |
| NP | 293 | R | R | R | R, ***K*** | K | K | Host-range (H1N1pdm)[9] |
| NP | 389 | K | ***R*** | R | R | R | R | Oligomer, T-cell epitope[10] |
| NA | 8 | I | I | I | I | I | ***T*** |  |
| NA | 16 | T | T | T | ***A*** | A | A |  |
| NA | 30 | A | A | A | A | A, ***T*** | T |  |
| NA | 50 | V | V | V | ***I*** | I | I |  |
| NA | 67 | Y | Y | Y | ***H*** | H | H, Y |  |
| NA | 147 | G | G, ***S*** | S | S | S | S | Oligomer, ligand binding (Oseltamivir) |
| NA | 153 | I | I, ***T*** | T | T | T | T | Oligomer, ligand binding (Oseltamivir) |
| NA | 155 | H | H | H | ***Y*** | Y | Y | Oligomer, ligand binding (Oseltamivir) |
| NA | 206 | V | V | V | V | V, ***I*** | I |  |
| NA | 263 | V | V, I | I | I | I, V | I | Oligomer, ligand binding (Oseltamivir) |
| NA | 283 | R | R | R | ***Q*** | Q | Q | Ligand binding |
| NA | 311 | S | S, ***N*** | N | N | N | N | Oligomer, ligand binding (Oseltamivir) |
| NA | 313 | D | D, ***N*** | N | N | N | N | Addition of N-glycan |
| NA | 328 | N | N | N | N | N, ***D*** | D | Oligomer, ligand binding (Oseltamivir), antigenic site |
| NA | 338 | R | R, ***K*** | K | K | K | K | Oligomer, ligand binding (Oseltamivir) |
| NA | 357 | E | E, ***D*** | D | D | D | D | Ligand binding |
| M1 | 15 | V | ***I*** | I | I | I | I |  |
| M1 | 95 | R | R | R | ***K*** | K | K | Oligomer |
| M1 | 207 | S | ***N*** | N | N | N | N |  |
| NS1 | 4 | N | N | N | N | N | ***K*** | Oligomer, ligand binding |
| NS1 | 47 | G, S | G, S | G | G | G | G |  |
| NS1 | 86 | A | ***V*** | V | V | V | V | Oligomer |
| NS1 | 197 | T | ***I*** | I | I | I | I | Oligomer, ligand binding |
| NS1 | 212 | P | P | P | ***S*** | S | S | Ligand binding[11] |
| NS1 | 218-331 | + | + | + | +, ***∆*** | ∆ | ∆ | Innate immune signaling[12] |

Table S3. Nonsynonymous fixed mutations to H3N2 CIV clades. Fixed mutations to H3N2 CIV lineage in bold/italics. Variant emergence in preceding clade noted as observed polymorphism among isolates. Variant fixation *within* clade is noted as polymorphism with bold/italics of latter fix. Note that Clade 3 is terminal (no subsequent lineages), and that Clade 4 also contains several terminal branches before establishment of Clade 5. HA = H3 numbering. Clear functions and references added when applicable.

| **Segment** | **Position** | **RCA** | **Variants** | **Lineage/Isolate** | **Notes/References** |
| --- | --- | --- | --- | --- | --- |
| PB2 | 38 | R | K | IL21 |  |
| PB2 | 76 | I | V | Los Angeles (CA) 21 |  |
| PB2 | 185 | I | V | TX23 |  |
| PB2 | 288 | Q | L | TX23 |  |
| PB2 | 315 | M | I | TX23 |  |
| PB2 | 340 | K | K, R | 2021 introduction |  |
| PB2 | 598 | A | A, T | 2021 introduction |  |
| PB2 | 676 | T | T, M | Los Angeles (CA) 21 |  |
| PB2 | 714 | I | T | Los Angeles (CA) 21 |  |
| PB1 | 215 | R | ***K*** | 2022 introduction |  |
| PB1 | 464 | D | ***N*** | 2022 introduction |  |
| PB1 | 581 | E | G | Los Angeles (CA) 21 |  |
| PB1 | 584 | R | H | TX21 |  |
| PB1 | 682 | I | V | Los Angeles (CA) 21 |  |
| PB1 | 688 | M | V | IL21 |  |
| PB1 | 744 | V | ***M*** | 2022 introduction |  |
| PA | 199 | E | K | FL21 |  |
| PA | 489 | C | S | FL21 |  |
| PA | 683 | L | ***F*** | 2022 introduction |  |
| HA1 | 43 | A | T | FL21 |  |
| HA1 | 91 | S | G | Los Angeles (CA) 21 |  |
| HA1 | 17 | N | ***T*** | 2022 introduction |  |
| HA1 | 223 | V | I | TX23 |  |
| HA1 | 245 | I | M | VA22 |  |
| HA2 | 113 | S | L |  |  |
| HA2 | 174 | K | ***R*** | 2022 introduction |  |
| NP | 105 | V | M | IL21 |  |
| NP | 189 | M | I | TN22 |  |
| NP | 402 | S | Y | Los Angeles (CA) 21 |  |
| NP | 430 | T | I | Los Angeles (CA) 21 |  |
| NP | 431 | G | E | Los Angeles (CA) 21 |  |
| NP | 446 | R | I | OK23 |  |
| NP | 485 | G | R | Los Angeles (CA) 21 |  |
| NA | 17 | I | L | TX21 |  |
| NA | 31 | T | I | Los Angeles (CA) 21 |  |
| NA | 53 | C | Y | TX22 |  |
| NA | 67 | H | ***Y*** | 2022 introduction |  |
| NA | 78 | C | F | TX22 |  |
| NA | 216 | G | S | Los Angeles (CA) 21 |  |
| NA | 248 | G | E | PA23 |  |
| NA | 332 | S | F | FL21 |  |
| M1 | 37 | T | I | Los Angeles (CA) 21 |  |
| M1 | 105 | R | ***K*** | 2022 introduction | Oligomer |
| NS1 | 37 | R | H | IL21 |  |
| NS1 | 59 | R | H | TX22 |  |
| NS1 | 89 | Y | H | IL21 |  |

Table S4. Nonsynonymous polymorphisms and fixed mutations in H3N2 CIV Clade 6 isolates. Relative to the recent common ancestor (RCA) residue. Fixed mutations found in all subsequent Clade 6 lineages are in bold/italics. Lineages correspond to geographic outbreak clusters identified in Figure 2. HA = H3 numbering. Clear functions and references added when applicable.

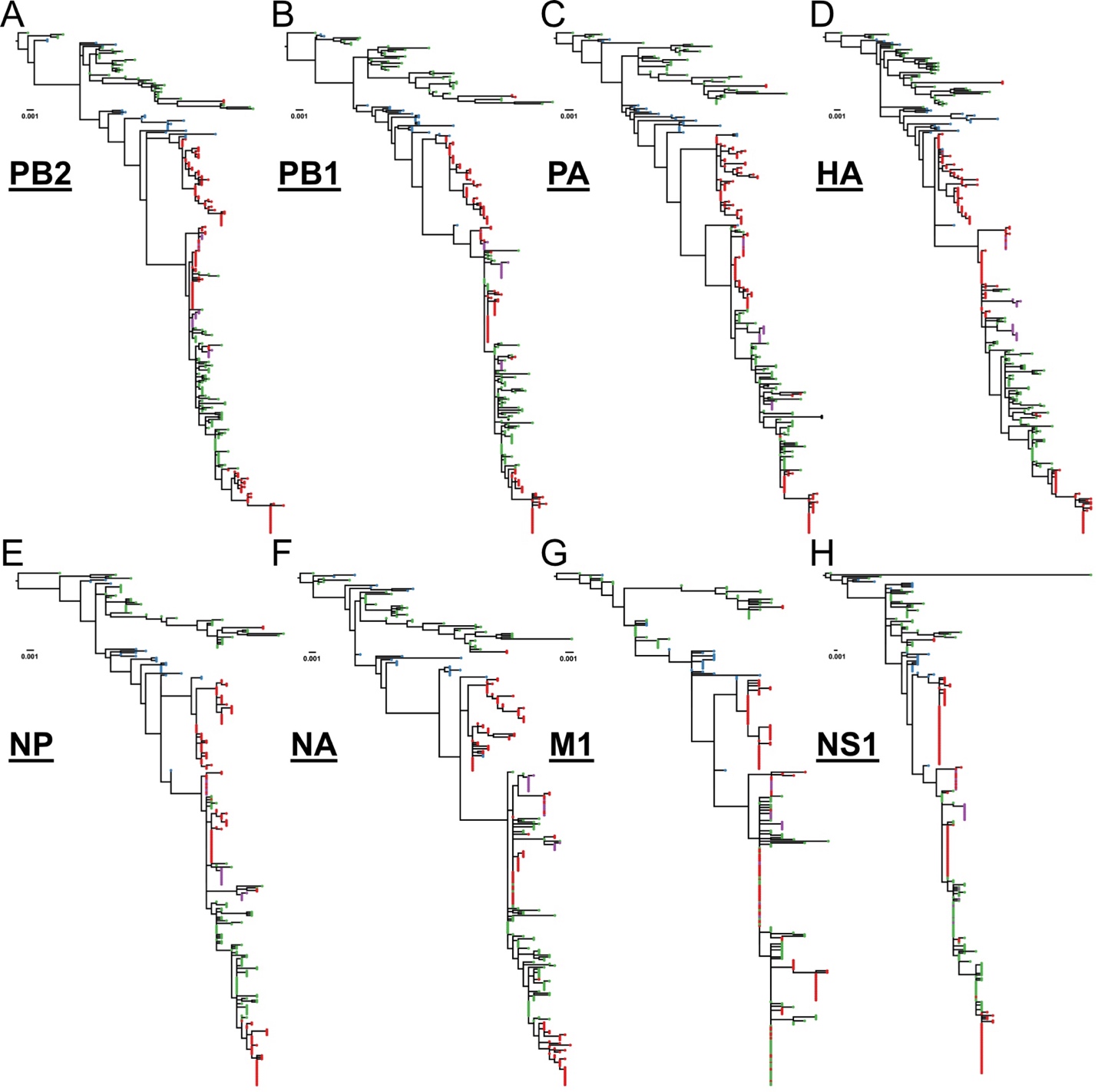

Figure S1. Maximum-likelihood (ML) trees of H3N2 CIV individual segment ORFs. Tips colored by country of sampling origin (green = China, blue = Korea, red = USA, purple = Canada).

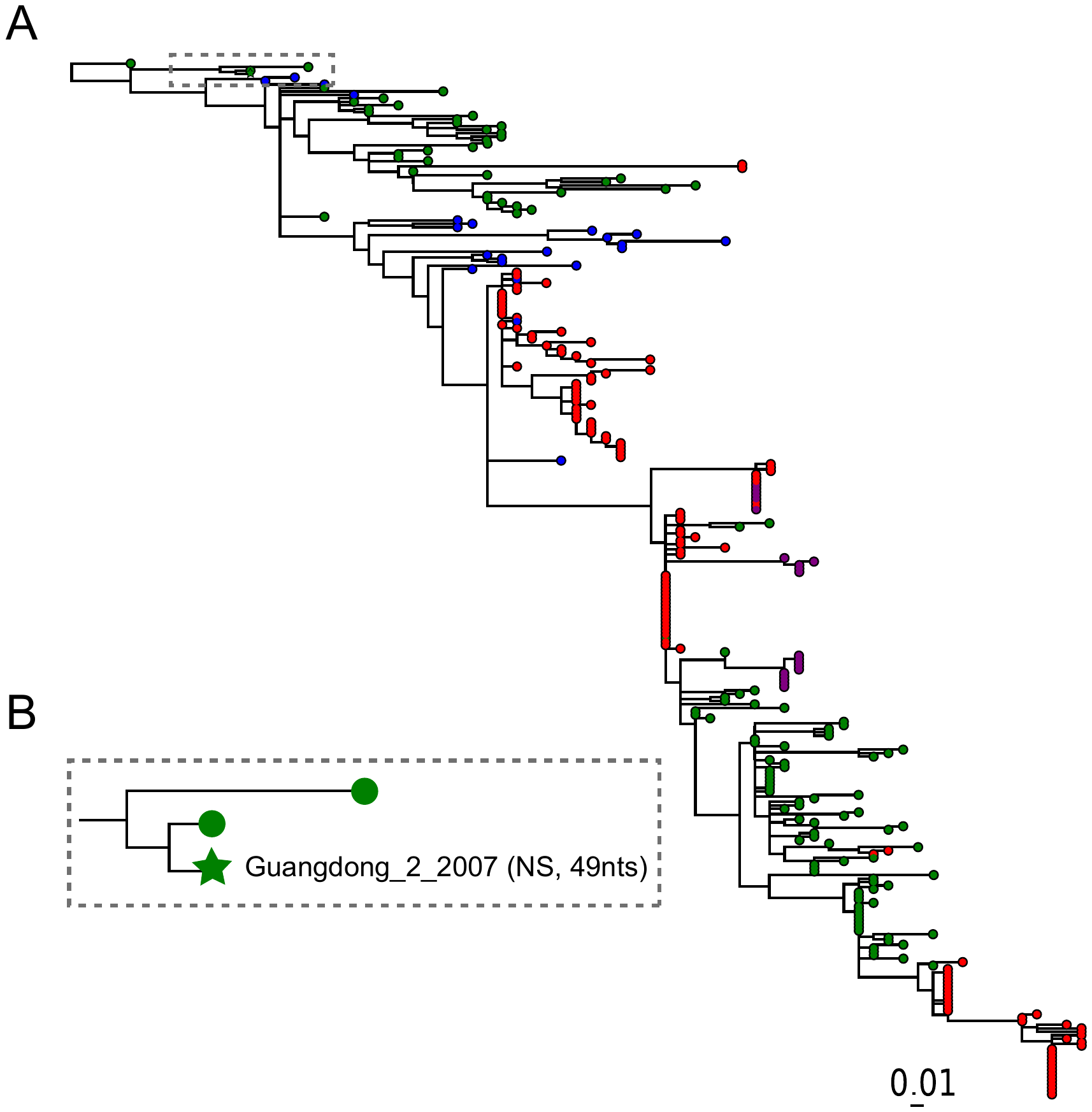

Figure S2. A single reassortment event was inferred for the NS gene in the Guangdong_2_2007 strain. (A) The TreeSort output for the 297 canine influenza sequences, using HA segment tree as the reference phylogeny. Tips colored by country of sampling origin (green = China, blue = Korea, red = USA, purple = Canada). The gray box highlights where on the phylogeny Treesort inferred a reassortment event, annotated with a star. (B) A zoomed in version of the inferred reassortment event. The reassorted NS gene segment for the Guangdong_2_2007 strain is estimated to differ by 49 nucleotide substitutions relative to the pre-reassorted segment.

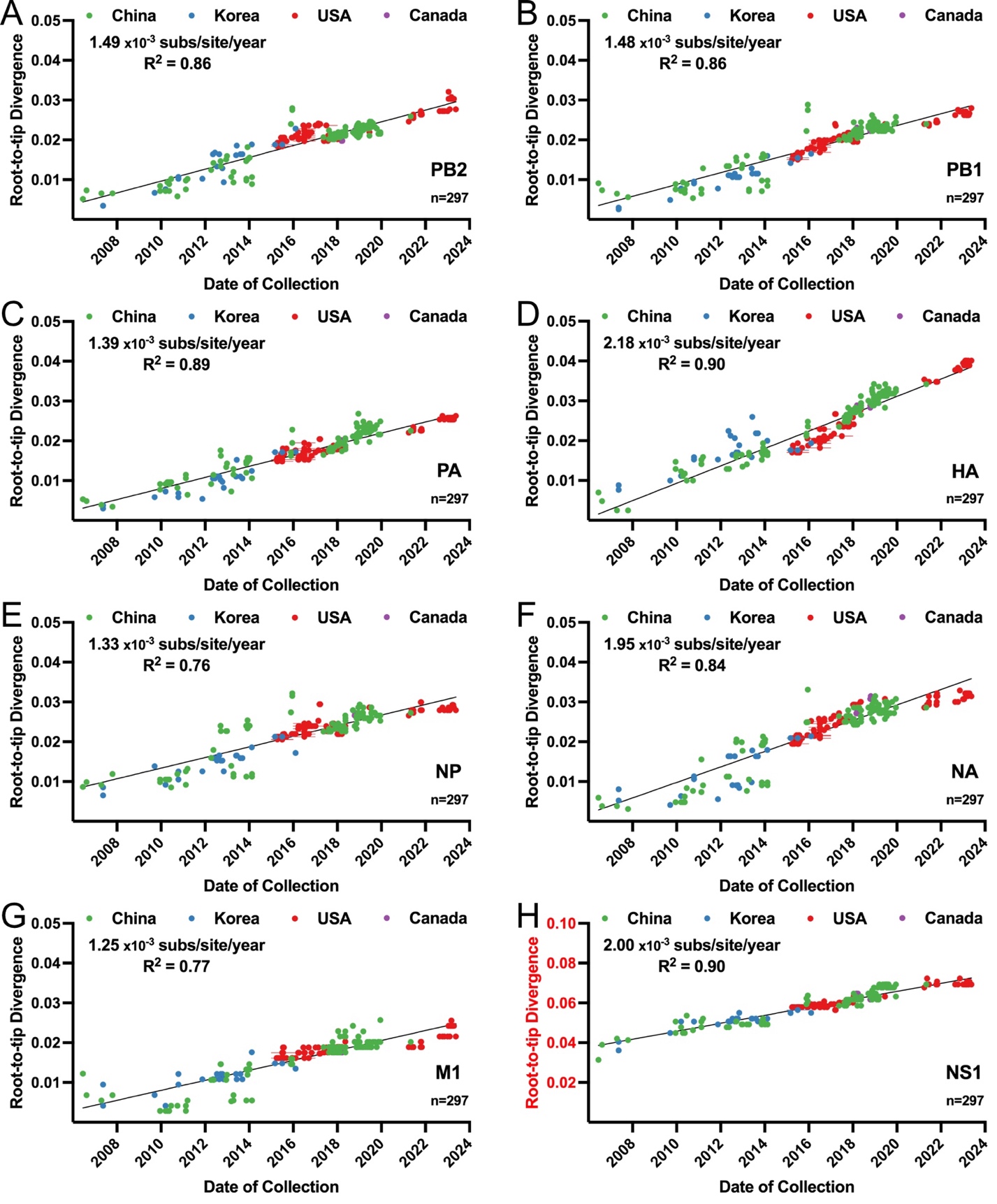

Figure S3. Root-to-tip divergence of individual CIV H3N2 segment ORFs. Calculated rates (corresponding to trendline) noted within figure. Note scale of divergence for NS1, resulting from a long root branch one of earliest samples (A/canine/Guangdong/1/2007) is a reassortant of this segment.

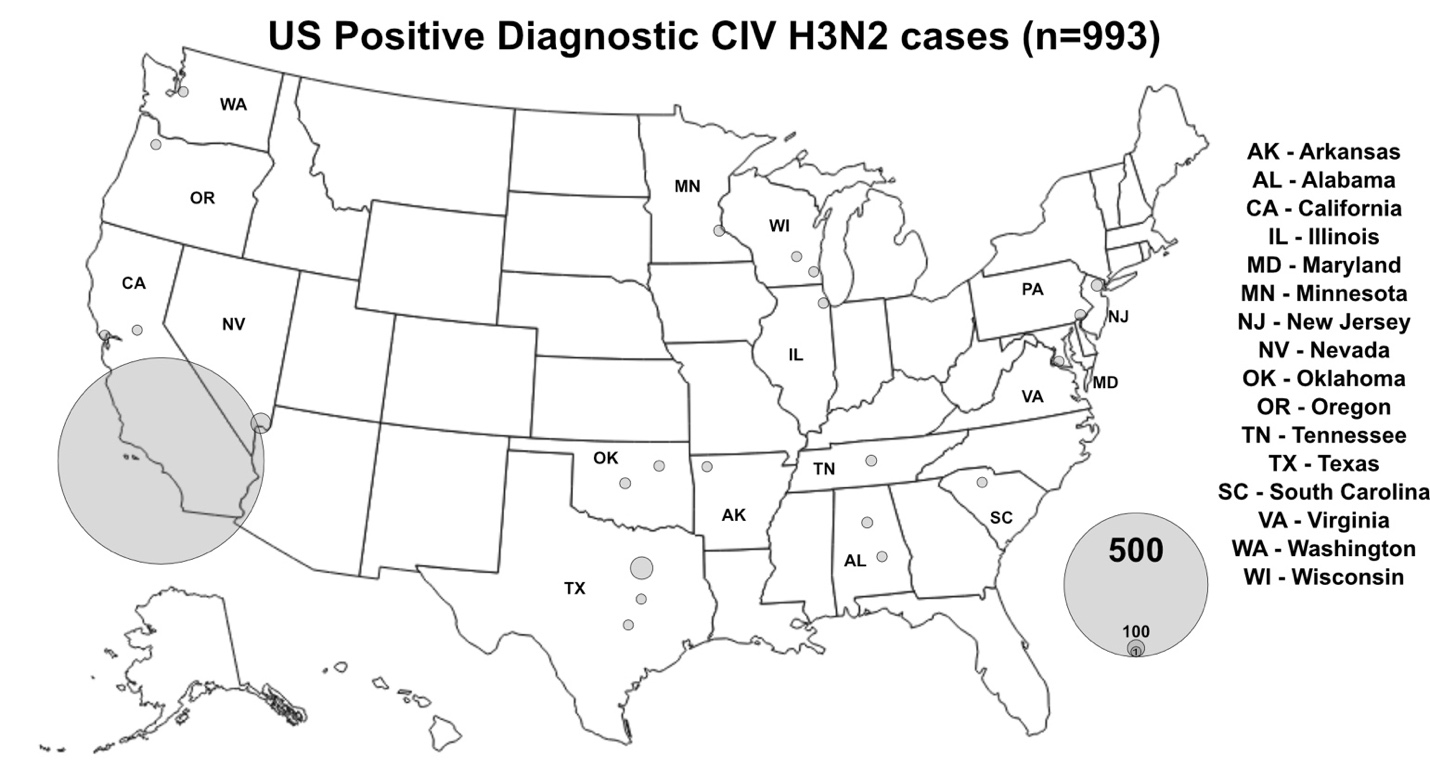

Figure S4. Diagnostic testing data set of CIV H3N2, (n=993). Logarithmic scale of cases.

Figure S5. MCC tree of common ancestor clade (n = 169) data set genomes of H3N2 CIV from Asia and North America from an analysis of discrete trait diffusion in BEAST v1.10.4. Color of tips and branches corresponds to the region of collection and inferred ancestral state reconstruction region respectively. The bars on nodes represent the 95% HPD on the node height.

**References – Supplemental**
